# Accurate de novo design of peptides from programming biophysical landscape

**DOI:** 10.64898/2026.06.08.731011

**Authors:** Mingyu Li, Yini Liu, Jie Zhong, Xinchao Shi, Jiqing Zheng, Kai Wang, Yunxia Cui, Chaoran Cheng, Shijin Li, Xiangzhe Kong, Shaoning Li, Miaojie Xv, Chunhao Zhu, Xiaobing Lan, He Yang, Kewei Chang, Zhenyu An, Shizhang Wan, Xiuyan Yang, Qiancheng Shen, H Eric Xu, Zihua Wang, Lei Liu, Youwen Zhuang, Jianzhu Ma, Jian Zhang

## Abstract

Peptides regulate virtually every cellular process and emerge as transformative modalities in bioengineering and therapeutics. However, the de novo design of functional peptides remains challenging, as peptide–protein interactions are governed by delicate and dynamic biophysics that are difficult to model, and experimental datasets are scarce. Here we present NeoPep, a generative deep-learning framework that encodes biophysical principles to accurately design functional peptides de novo. By integrating over 5 million peptide–protein complexes spanning experimentally determined, sequence mimics, and structure ensembles, NeoPep learns the complex biophysical landscape governing peptide function and enables its programmable control. In prospective application across 10 diverse and challenging targets, NeoPep generated potent peptide binders, agonists, and antagonists with hit rates of 12.5–66.7%, even in the absence of defined binding sites or structure information. Beyond de novo co-design, NeoPep supports standalone structure or sequence redesign, readily discriminating subtle context differences. In the structure redesign mode, it generates highly selective peptides with atomic-level conformational accuracy (Cα RMSD < 2.0 Å to our solved cryo-EM structure). This structural precision circumvents the need for experimental structure determination, further accelerating iterative sequence redesign to yield a 43.3-fold improvement in potency. These findings establish a general framework for translating biophysical principles into actionable peptide functions, with broad implications for basic research, bioengineering, and medicine.

## Introduction

Peptides are pervasive biological regulators whose short sequences can readily evolve to engage diverse molecular targets through structural and functional plasticity^1,2^. Occupying a size niche between small molecules and large biologics, peptides combine the favorable properties of both modalities and have emerged as a promising frontier for therapeutic development and bioengineering applications^3–5^. Motivated by this goldilocks regime, considerable effort has been devoted to the discovery of bioactive peptides through high-throughput screening platforms, including display- and library-based techniques^6^, as well as natural product mining^7^ (Fig. 1a). While powerful, these approaches largely rely on empirical exploration of vast chemical spaces, making them costly, labor-intensive, and difficult to direct toward specific binding modes for optimization.

**Fig. 1.**
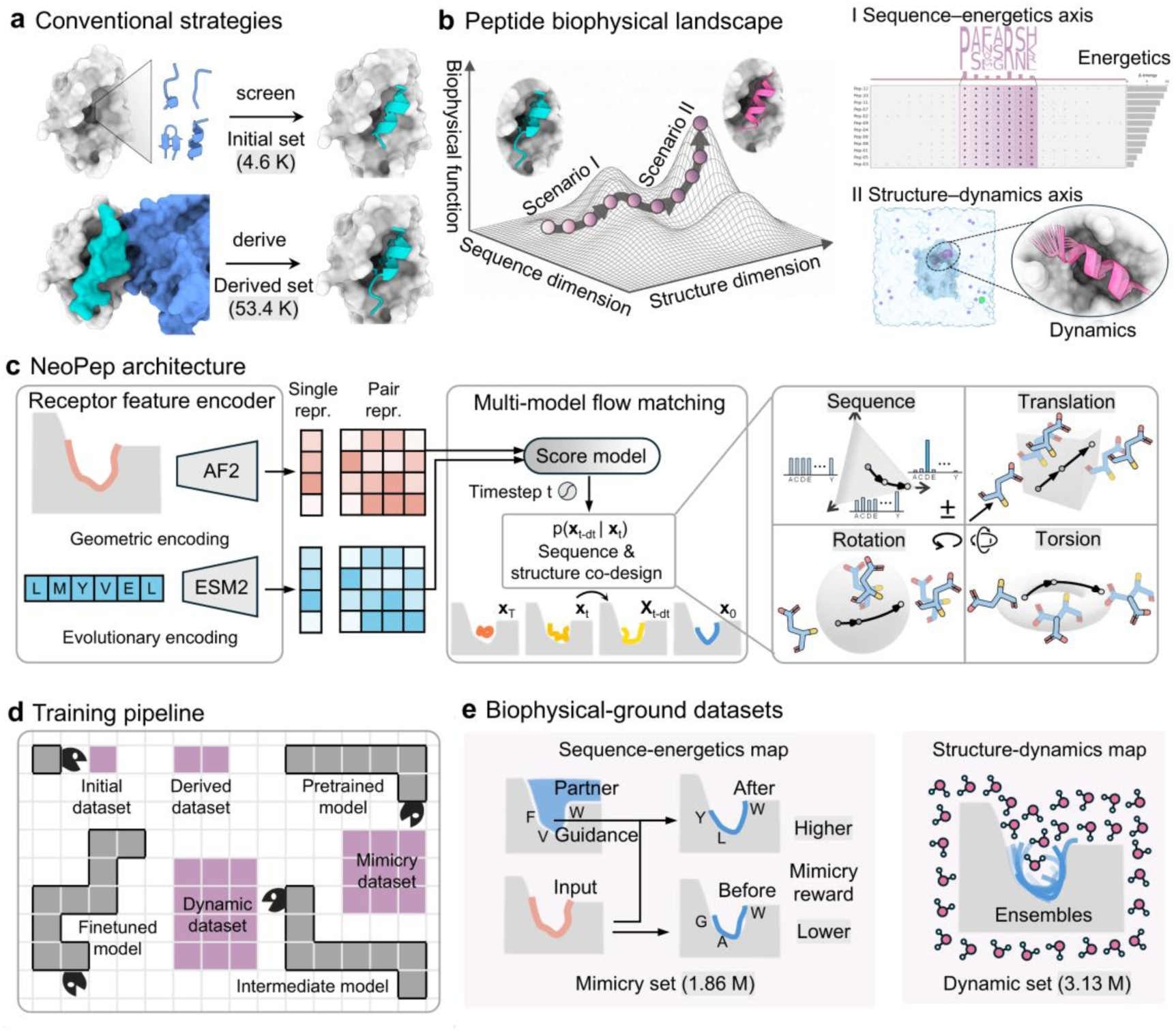
Overview of NeoPep model. **a,** Conventional peptide design strategies. **b,** Biophysical landscape of peptide binding. Integrating molecular modelling data enables learning of a biophysics-informed sequence–structure landscape to guide peptide optimization. **c,** NeoPep architecture consisting of receptor feature encoder, score model and multi-modal flow matching framework. **d,** Model training process, conceptualized as a Snake game. **e,** Biophysical-ground dataset curation.

Computational design offers a compelling alternative. Although easy-to-use platforms have revolutionized protein design^8–14^, equivalent tools for peptide design remain elusive, hindered by the distinct and challenging sequence–structure biophysical landscape that governs peptide function. Longer proteins leverage cooperative long-range interactions to stabilize compact, ordered secondary structures and/or tertiary folds. Short peptides, by contrast, are dictated predominantly by local sequence propensities and short-range interactions, frequently sampling disordered coil states unless pre-organized by helical or cyclically constrained topologies to reduce entropic penalties^15–17^. Consequently, despite progress in zero-shot protein design models^18–24^, they tend to explore only a narrow region of peptide design landscape, favoring longer, less flexible helical binders or conformationally constrained peptides. Moreover, short peptides exhibit weak alignment-detectable evolutionary conservation and present small, delicate binding interface, such that even minor geometric inaccuracies during modeling may incur amplified energetic penalties^25^.

The intrinsic dynamics of peptide–protein interactions further hamper conventional structural determination method by X-ray crystallography and cryo-electron microscopy (cryo-EM). As a result, peptide–protein complexes account for merely ∼3.5–5% of Protein Data Bank (PDB) entries, roughly a quarter of the available small-molecule complexes and an order of magnitude fewer than protein–protein complexes. Moreover, these experimental structures typically represent static snapshots that fail to capture the broader conformational ensembles underlying peptide function. Such paucity of training data imposes a few-shot learning constraint on protein language models (pLMs) or structural foundation models^26–29^, limiting their capacity to map the peptide biophysical landscape despite recent progress. Importantly, their utility has not yet been rigorously assessed in real-world de novo peptide design across a wide array of protein targets.

These barriers underscore the critical need for large-scale datasets that explicitly decipher the localized atomistic interactions and profound conformational heterogeneity inherent to peptides. To map this complexity, we decompose the biophysical landscape of peptide function into two tractable axes (Fig. 1b): sequence–energetics, focusing on how sequence complementarity drives interaction energy, and structure–dynamics, capturing how structural plasticity mediates functional diversity. Molecular modelling, grounded in established biophysical principles and decades of experimental validation, provides a versatile tool for probing both axes of peptide behavior^30–32^. We reasoned that such biophysics-based modelling could generate the large-scale datasets necessary to extend generative foundation models with genuine biophysical awareness.

Here, we developed a neopeptide designer called NeoPep, a generative deep-learning engine that programs the biophysical landscapes to deliver accurate peptide engineering. NeoPep is built upon three core pillars: (i) it empowers structure foundation models with contextualized sequence representations from pLMs, trained jointly on a curated dataset derived from over 1 million protein-protein interactions (PPIs). (ii) it employs a reinforcement-based, self-distilled extension coupled with molecular modeling, generating 1.86 million energetically favorable sequence variants to fine-tune the sequence–energetics axis. (iii) it explores the structure–dynamics axis by fine-tuning on 313 μs molecular dynamics (MD) conformational ensembles spanning 3.13 million conformations. In computational benchmarks, NeoPep-generated peptides exhibit high binding fitness, matching or exceeding those of experimentally verified complexes and outperforming the state-of-the-art RFdiffusion model in targeting structured proteins with peptides. In experimental validation across a diverse set of 10 challenging protein targets, NeoPep achieves hit rates of 12.5–66.7%, identifying potent binders, antagonists, and agonists. Beyond sequence-structure co-design, NeoPep supports independent structure prediction and sequence optimization, enabling the discovery of the signaling-selective peptides through structure resampling, and the 43.3-fold potent peptide via iterative sequence redesign.

## Main

### Design of the NeoPep model

Unlike general-purpose protein design platforms, NeoPep is specifically engineered for short peptides. It builds on an architecture that harnesses the synergistic capabilities of sequence and structure foundation models: to incorporate rich evolutionary signals from the pretrained ESM2 pLMs^14^ and to construct precise geometric interactions from the trainable AlphaFold2 (AF2) invariant point attention (IPA) transformers^10^. A novel fusion module scales and projects the single and pair representations from both input modalities into a shared latent space, which feeds into a multi-flow matching score model that jointly generates target-binding peptide sequence and structure (Fig. 1c and Materials and methods).

A central obstacle in training peptide-specific models is the absence of high-quality data disclosing the peptide biophysical landscape. To this end, NeoPep was trained through a three-stage curriculum, integrating curated data from diverse, complementary sources (Fig. 1d, Supplementary Fig. 1 and Materials and methods). Firstly, NeoPep was pretrained on a combined dataset, comprising an initial set of experimentally determined peptide–receptor complexes and a derived set extracted from million-scale PPIs (Technical details are provided in the “Curation of training datasets” section in the Supplementary Methods). Briefly, recognizing that a substantial proportion of PPIs are driven by peptide-like hotspot segments that contribute the majority of the binding energy^25^, we applied the established Rosetta PeptiDerive protocol^33^, a computational analogue of peptide microarray screening, to scan and derive these segments. This procedure yielded an order-of-magnitude more data pairs than the Initial dataset.

To further capture the biophysical landscape, the subsequent fine-tuning proceeded along two complementary axes. Along the sequence axis, we fine-tuned the model on 1.86 million sequence variants from a Mimicry dataset (Fig. 1e, left). Peptide mimicry is a commonly used strategy wherein candidates are designed to preserve or enhance the activity of known bioactive peptides or protein interaction motifs by mimicking their key interfacial interactions^3,34^. To instill mimicry capacity, we developed a training-free guided algorithm that efficiently steers the pretrained model toward higher interface mimicry reward while maintaining proximity to the prior learned distribution (Supplementary Fig. 2 and evaluations are provided in the “Details of mimicry-guided flow matching framework” section of the Supplementary Methods). A subsequent self-distillation strategy was carried out: the pretrained, mimicry-guided model generated >8 million peptide mimics targeting the receptors from the Initial and Derived datasets. The final high-quality Mimicry dataset was obtained by meticulously filtering these candidates using biophysics-based criteria computed from Rosetta molecular modeling. Along the structure axis, we generated all-atom MD trajectories in explicit solvents, resulting in a Dynamic dataset accumulating over 313 μs and 3.13 million MD conformation data (Fig. 1e, right). Analysis of key thermodynamic properties confirmed equilibration across the simulations and the consistently low dissociation rate of the Derived dataset supports the validity of the extracted peptide hotspot segments (Supplementary Fig. 3). The model was then fine-tuned on the Dynamic dataset to embed these conformational ensembles. Such sequential fine-tuning order also serves to mitigate the risk of the model overfitting to the self-distilled, noisy sequence variants.

### In silico analysis of NeoPep designs

The primary objective of target-conditioned generative methods is to maximize binding fitness by producing peptides with high-affinity sequence variants and feasible bound conformations. We assessed binding affinity using Rosetta interface Gibbs free energy change (ΔG) upon binding and conformational feasibility using the Cα root mean square deviation (RMSD) and all-atom interface DockQ scores relative to the native binders. The performance of NeoPep is compared against baselines, ranging from the autoregressive HSRN^35^ and iterative non-autoregressive dyMEAN^36^ to the diffusion-based RFDiffusion^11^ (Supplementary Methods).

At the pretraining stage, NeoPep outperformed all baselines in binding fitness, generating peptide binders with higher affinities and feasible native-like conformations. The ΔG distribution of NeoPep-derived peptide (−30.3 kcal/mol) most closely approximated that of native peptides (−36.3 kcal/mol) (Fig. 2a). Meanwhile, NeoPep showed a median Cα RMSD of 2.6 Å with 47.3 % of cases below the 2.5 Å success threshold (Fig. 2b), and a DockQ score of 0.72 with 44.1% above the 0.80 high quality threshold (Fig. 2c), whereas the second-best baseline reached 7.22 Å Cα RMSD (one-sided paired t test, *p* = 2.5×10^−16^) and 0.40 DockQ (one-sided paired t test, *p* = 1.7×10^−28^). Compared to PepFlow^27^, NeoPep demonstrated substantially improved binding affinity (−5.5 kcal/mol, one-sided paired t test, *p* = 5.2×10^−3^) with marginal improvement in binding conformation (DockQ, one-sided paired t test, *p* = 5.6×10^−5^). This suggests that the added benefit of ESM2-derived evolutionary priors primarily expanding sequence space to optimize affinity.

**Fig. 2.**
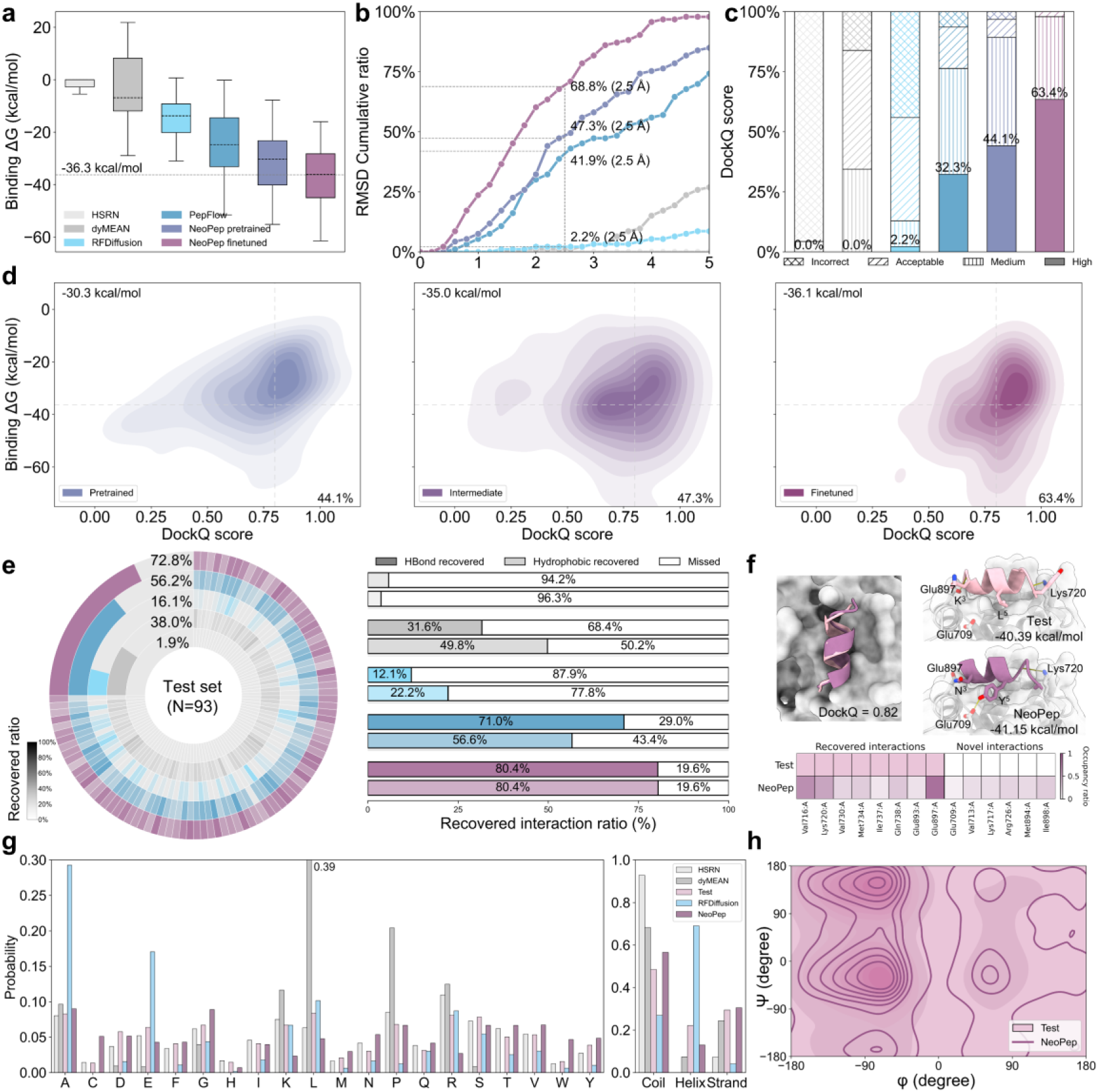
In silico benchmarking of de novo target-binding peptide generation. **a,** Distribution of Rosetta ΔG. The dashed line indicates the median binding ΔG of native test peptides. Box plots show the median (center line), interquartile range (IQR; box limits), and 1.5× IQR (whiskers). The color scheme applies to all subsequent panels.**b,** Cumulative frequency of peptide Cα RMSD. The dashed line denotes the success rate for high-accuracy structural recovery (RMSD ≤ 2.5 Å). **c,** Stacked histogram of all-atom interface DockQ scores. The proportion of cases meeting the high-quality threshold is annotated. **d,** Joint density of binding ΔG versus DockQ across NeoPep curriculum learning stages. Median ΔG are annotated on the left while percentages on the right indicate the fraction of high-quality DockQ predictions. **e,** Recovery of native interactions. Left, interaction recovery ratio per case, with darker shades indicating higher recovery and the overall recovery ratio shown in the second quadrant. Right, Differential recovery of hydrogen bonds (HBonds) and hydrophobic contacts. **f,** Structural superposition of experimentally determined (pink, SEKFKLLFQSY) and NeoPep-designed (purple, VSNLEVYRSQA) peptides in complex with target receptors (gray). Key interacting residues are shown as sticks. Bottom panels detail interaction statistics relative to the native binder, where color intensity correlates with generation frequency. **g,** Probability distributions of amino acid composition and secondary structure elements for generated peptides across models against the native reference (pink). **h,** Ramachandran plot comparing backbone dihedral angles (φ and ψ). The native distribution (pink) is overlaid with density contours of the NeoPep-generated (purple).

Sequential fine-tuning on both sequence and structure dimensions extended the model beyond pretraining to encapsulate a broader biophysical landscape. Mapping the joint distribution of binding ΔG versus DockQ across three training stages uncovered a global shift toward the lower-right quadrant, signifying progressively higher binding fitness (Fig. 2d). The intermediate model yielded a substantial gain in binding ΔG (−4.7 kcal/mol, from −30.3 to −35.0 kcal/mol), whereas the fine-tuned model exhibited a marginal improvement of −1.1 kcal/mol (to −36.1 kcal/mol). This pattern, characterized by energy-dominant gains after explicitly fine-tuning on millions of sequence variants, is consistent with the improved pretraining performance relative to PepFlow after implicitly incorporating ESM2-derived sequence priors. On the contrary, the intermediate model presented only a slight binding Cα RMSD improvement (0.3 Å, from 2.6 to 2.3 Å), while the fine-tuned model exhibited an improvement of 0.6 Å, reaching a high-resolution RMSD of 1.7 Å. Meanwhile, the DockQ distribution shifted from a bimodal pattern to a single high-quality peak. Under stringent criterion (DockQ > 0.80), the intermediate and fine-tuned models increased the proportion of high-quality designs by 3.2% and 16.1%, respectively, culminating in 63.4% of generated complexes adopting high-quality conformation.

The robustness of fine-tuned model (hereafter, “NeoPep” refers to the fine-tuned model unless otherwise specified) over existing baselines are further evaluated across both intra-peptide properties, including length and secondary structure, and inter-binding pocket depth (Supplementary Fig. 4). From an intra-molecular perspective, NeoPep achieves the most stable binding ΔG across all peptide length. It is observed that ΔG generally decreased as peptide length increased, primarily due to the formation of more extensive interface areas (Supplementary Fig. 5). Conversely, DockQ scores exhibited an inverse trend, declining for longer peptides mainly due to their expanded conformational search space. NeoPep, nevertheless, demonstrates superior robustness to sequence length, maintaining a near-high average quality (DockQ > 0.66), while baselines experience a sharp decline to incorrect quality level (DockQ < 0.23). Moreover, secondary structure categories appear to have minimal impact on NeoPep’s binding fitness. From an inter-molecular perspective, PepFlow degrades from high to acceptable quality (DockQ < 0.49) in shallow pockets, while RFDiffusion and dyMEAN struggle between acceptable and incorrect quality. In contrast, NeoPep remained largely unaffected by pocket geometry (DockQ > 0.69).

Beyond confirming binder viability, a deeper examination was conducted to assess the residue-level interactions that drive binding fitness and functional modulation (Fig. 2e). Using the PLIP to map inter-molecular interactions across native and generated complexes^37^, we found that NeoPep recovered a leading 72.8% of native contacts. In addition, 51.0% of the contacts by NeoPep-designed peptides were newly formed, suggesting that NeoPep not only reliably recapitulates key native contact patterns but also actively explores alternative productive interactions. In-depth analysis of the recovered interaction types revealed that RFDiffusion and dyMEAN preferentially recovered hydrophobic contacts, whereas PepFlow favored hydrogen-bond recovery. NeoPep maintained a balanced profile by recovering 80.4% of both hydrogen and hydrophobic pairs. To illustrate these capabilities, we analyzed an exemplary peptide designed for the androgen receptor (PDB ID: 1T5Z) from the test set (Fig. 2f). NeoPep generated a sequence distinct from the native peptide while preserving a near-native binding mode (DockQ = 0.82). Interaction analysis revealed that the design retained two critical hydrogen bonds: one utilizing the same backbone atom to engage K720, and another where the substituted N^3^ sidechain replaced the native K^3^ backbone interaction with E897. Moreover, the L^10^Y mutation introduces a hydroxyl group that extends hydrogen bonding to E709. Together, these favorable polar interactions provide a structural basis for the enhanced binding affinity (ΔΔG = −0.76 kcal/mol) of the designed peptide relative to the native reference.

It is essential to validate generated peptides by assessing their consistency with realistic sequence and structural distributions. We quantified the divergence between generated and test sets across three metrics: primary amino acid composition, secondary structure distributions and backbone amide Ramachandran angles. First, the amino acid distribution produced by NeoPep aligns more closely with test set, exhibiting a Kullback–Leibler (KL) divergence of 0.13. In comparison, RFDiffusion (KL = 0.38) and dyMEAN (KL = 0.89) exhibit strong biases towards alanine (A), glutamate (E), leucine (L), and proline (P) (Fig. 2g). Second, NeoPep shows the highest fidelity to the native secondary structure distributions, with coil, helix and strand proportions of 0.56, 0.13 and 0.30, respectively, deviating from the test set by less than 0.10. As we introduced, RFDiffusion displays a strong propensity for helical geometries (0.69), which likely reflects its overrepresentation of the highly helix-promoting alanine residue^38^ (Fig. 2g). Third, the Ramachandran plots uncovered that NeoPep accurately recapitulates native backbone dihedral distributions, with sampled conformations concentrated in the favorable upper-left and bottom-left quadrants (Fig. 2h).

### Experimental testing of de novo peptides

After curriculum learning on the sequence Mimicry and structure Dynamic datasets, *in silico* evaluations demonstrate that NeoPep broadly samples the sequence axis while providing high-quality initial binding poses. Nevertheless, recognizing such pose is incomplete for intrinsic flexible peptides, we deploy explicit-solvent MD simulations to expand this initial seed into the broader conformational ensembles, then quantify their thermodynamic stability through ensemble-averaged molecular mechanics Poisson−Boltzmann surface area (MM/PBSA) binding free energies^39^ (Fig. 3a, Supplementary Methods). For comparison, the initial seeds were re-evaluated using AF2-multimer^8^ and also validated via MD. Following sequence similarity reduction, 48 candidate peptides comprising 24 NeoPep-designed and 24 AF2-multimer redocked were synthesized for bioassays (Materials and methods).

**Fig. 3.**
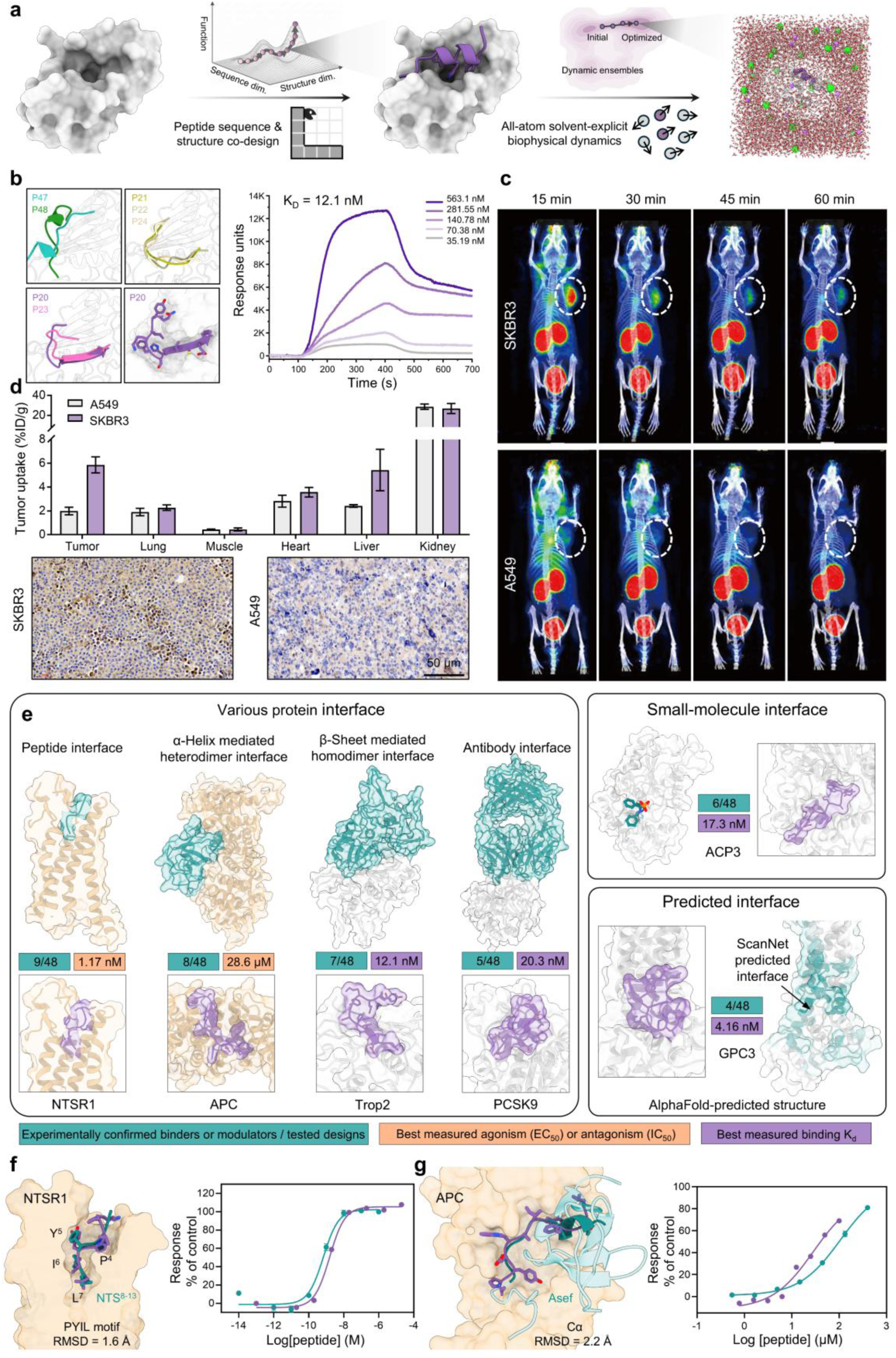
Experimental verification of de novo peptides designed by NeoPep. **a,** Schematic of the NeoPep pipeline for peptide sequence and structure co-design. **b,** Structural and affinity characterization of identified Trop2-binding peptides. Left, Trop2 (light grey) complexed with validated binders adopting partially helical, β-sheet, or loop conformations. Right, SPRi sensorgrams of Trop2-P20 against Trop2. **c,** Micro-PET/CT images at various time points after intravenous injection of peptides in SKBR3 and A549 tumor-bearing mice (n = 3). Tumors are indicated by white circles. **d,** Tumor uptake (%ID/g) in SKBR3 and A549 models (left), alongside immunohistochemistry staining of Trop2 expression in corresponding tumor tissue sections (right). Scale bar, 50 μm. Data are represented as mean ± SEM (n = 3). **e,** Overview of protein targets for peptide design. Target surfaces are colored wheat or grey, with native binders in green and NeoPep-designed peptides in purple. Green boxes indicate the number of observed hits (binders or modulators) out of total tested designs. Purple and orange boxes show the K_d_ of the highest-affinity binder and the IC_50_/EC_50_ of the most potent functional modulator, respectively. **f, g,** Generated structures and functional validation of designed peptides targeting NTSR1 (**f**) and APC (**g**). Left, target receptors (wheat surface) bound to the native peptide (NTS^8–13^ for NTSR1) or protein partner (Asef for APC) in blue, overlaid with NeoPep-designed peptides in purple. Right, dose–response curves comparing the native or derived peptide (blue) against NeoPep-designed peptides (purple). Data are mean ± SEM (n = 3).

We first validated the pipeline by generating peptides against the homodimer interface of Trop2 and verified binding affinity by surface plasmon resonance imaging (SPRi). Five NeoPep-derived peptides bound Trop2 with dissociation constants (K_D_) ranging from 12.1 nM (Trop2-P20) to 2.2 μM, while only two AF2-multimer redocked peptides exhibited weaker K_D_ values of 0.54 μM and 7.3 μM. Structural modeling uncovers diverse generated conformations, with the highest-affinity peptide, Trop2-P20, adopting a hairpin-like pose tightly anchored within the β-sheet mediated interface (Fig. 3b). As an empirical benchmark against conventional screening methods, a random decapeptide library with a sequence diversity of 10^7^ was constructed and screened against Trop2 (Supplementary methods). A total of 41 peptides were obtained with K_D_ ∼ 1 μM, implying that NeoPep delivers a 50,813-fold improvement in hit rate compared to experimental screening for this target. To decouple the contribution of post-MD conformational sampling, an independent set of 24 peptides was evaluated without this step, and two still reached K_D_ ∼ 1 μM, underscoring NeoPep’s capacity to generate reliable binders from scratch (Supplementary Table 1).

To evaluate *in vivo* targeting, the peptide P20 was conjugated with 1,4,7,10-tetraazacyclododecane-1,4,7,10-tetraacetic acid (DOTA) for chelation of gallium-68 and administered by tail-vein injection at 50 μM to mice bearing Trop2-positive SKBR3 tumors or Trop2-negative A549 tumors (Fig. 3c, Materials and methods). The imaging results from positron emission tomography (PET)–computed tomography (CT) uncovered a significant, 3.0-fold greater uptake in the SKBR3 tumors compared to the A549 tumors (one-sided unpaired t test, p = 3.2×10^−3^). Over time, the majority of the probe cleared renally within 60 minutes, with higher kidney signals reflecting expected metabolic accumulation. In accordance with the PET imaging, immunohistochemistry staining of tumor tissues confirmed that the designed peptide specifically recognized the Trop2-positive tumors with negligible off-target binding in the A549 cohort (Fig. 3d, Materials and methods).

Building on these results, we conducted a systematic experimental campaign across a diverse array of therapeutic targets and interface classes (Fig. 3e, right). Alongside the common protein-mediated interface of Trop2, we targeted the respective antibody, small-molecule, and ScanNet-detected^40^ interfaces of PCSK9, ACP3, and GPC3. NeoPep consistently identified three peptide hits for each, whereas AF2-multimer redocking yielded fewer hits across the same targets (two, three, and one hit, respectively). Across all four interface classes, the top NeoPep candidates achieved binding affinities in single- to double-digit nanomolar range (Supplementary Table 1).

Beyond binder design, we investigated whether NeoPep-designed peptides could act as functional modulators across varied topologies, extending from the β-sheet-mediated Trop2 interface to the NTSR1 peptide-binding site for agonism and the α-helix-dominated APC interface for antagonism (Fig. 3e, left). For NTSR1, six of 24 NeoPep-derived candidates triggered potent agonism, with the lead displaying an EC_50_ of 1.2 nM, comparable to that of the endogenous ligand NTS^8–13^ (0.70 nM). By comparison, AF2-multimer redocking identified three functional agonists with weaker potency (10.3 nM) (see the “Assessment of functional motif recognition for NTSR1” section of the Supplementary Methods). Structure analysis confirmed that NeoPep accurately captures functional binding topologies rather than mere sequence patterns (PYIL motif RMSD = 1.6 Å), which may account for its higher success rate (Fig. 3f). For APC, previous efforts systematically synthesized a series of truncated hotspot segments (5 to 19 residues) derived from the APC-binding region of Asef^41^. While the optimal 19-mer achieved an IC_50_ of 51.0 μM, most shorter segments exhibited IC_50_ above 100 μM. NeoPep produced five distinct de novo antagonists, four of which showed IC_50_ at or below 50 μM. The top candidate, a 12-mer (APC-P20; 28.6 μM), outperformed a comparably sized Asef-derived 13-mer (IC_50_ = 106.3 μM) by 3.7-fold. Instead, AF2-multimer redocking produced three weaker candidates (50.8 to 176.7 μM). Interestingly, despite peptide sequence divergence and the strict exclusion of homologous receptors, the binding orientation of APC-P20 closely mirrored that of the Asef binding hotspot segment (Cα RMSD = 2.2 Å) (Fig. 3g and Supplementary Table 1).

A retrospective analysis was undertaken to determine which *in silico* metrics most reliably predicted experimental success (Supplementary Fig. 6a). Across the diverse set of six targets, dynamic MM/PBSA ΔG derived from MD simulations successfully discriminated active from inactive candidates in three of six cases. Notably, this overall success rate conceals a pronounced disparity in performance between the input methods: discrimination was achieved in five of six NeoPep-generated candidate sets, compared with only one of six AF2-multimer redocked sets. Other evaluated metrics, spanning sequence length, secondary structure, physicochemical properties (hydrophobicity, isoelectric point, and charge at pH 7.0), static Rosetta ΔG, and AF2-multimer confidence scores (pLDDT and ipTM), demonstrated no meaningful predictive ability. To further evaluate sequence diversity, we projected the NeoPep-identified hits onto the sequence space sampled from the Initial and Derived datasets (Supplementary Fig. 6b). These hits exhibited a broad and uniform distribution throughout this space, indicating coverage across diverse sequence motifs.

### Access to non-canonical binding sites

Having established NeoPep’s reliability for bioactive peptide discovery across a broad spectrum of canonical protein targets, we next challenged the framework with two under-explored systems characterized by low data availability and intractable experimental structures. We directed our efforts toward the SIRT3 allosteric site and the GPR75 orphan receptor. These two non-canonical targets lack established peptides and experimental structures, yet hold compelling therapeutic opportunities.

SIRT3, one of the seven sirtuin family members, deacetylates acetylated peptide substrates at its orthosteric site^42^. It has been deemed difficult to drug because its high-affinity endogenous substrates preclude competitive inhibition, and the orthosteric pocket is not amenable to activation. Allosteric regulation from a topologically distant site offers a promising alternative, conferring subtle upregulation or downregulation of enzymatic activity^43^ (Fig. 4a). We had previously identified a spacious allosteric site in SIRT3 using a macrocyclic sulfonamide agonist (EC_50_ = 29.4 μM)^44^. Here, we utilized this site as the structural input for NeoPep. Of note, AF2-multimer redocking consistently predicted all 1,000 peptides to bind the orthosteric site. Such bias is plausible, given that numerous solved sirtuin structures in its training data normally contain a bound orthosteric peptide substrate (Supplementary Fig. 7). Nevertheless, we synthesized the top 24 NeoPep-generated and 24 AF2-multimer redocked peptides for comparison. Initial screening detected six peptides exhibiting either antagonistic or agonistic activity (Fig. 4b). For antagonism, the AF2-multimer redocked SIRT3-P25 was more potent (IC_50_ = 1.5 μM) than the NeoPep-direct SIRT3-P1 (IC_50_ = 36.5 μM). For agonism, NeoPep-derived candidates excelled: SIRT3-P24 was the most potent agonist (EC_50_ = 1.0 μM) and SIRT3-P22 produced the highest efficacy (span = 2.2-fold) (Fig. 4c and Supplementary Table 2). Structural modeling of the representative NeoPep-derived antagonist and agonist revealed characteristic L-shaped engagement of the allosteric pocket, with a conserved C-terminal phenylalanine anchoring each peptide while the flexible body adopts distinct conformations to exert differential regulation (Fig. 4d, left). The observed agonism of SIRT3-P48, however marginal, directly contradicts its AF2-multimer predicted binding pose. The complete spatial overlap would inherently displace the substrate and thus drive competitive inhibition rather than activation (Fig. 4d, right).

**Fig. 4.**
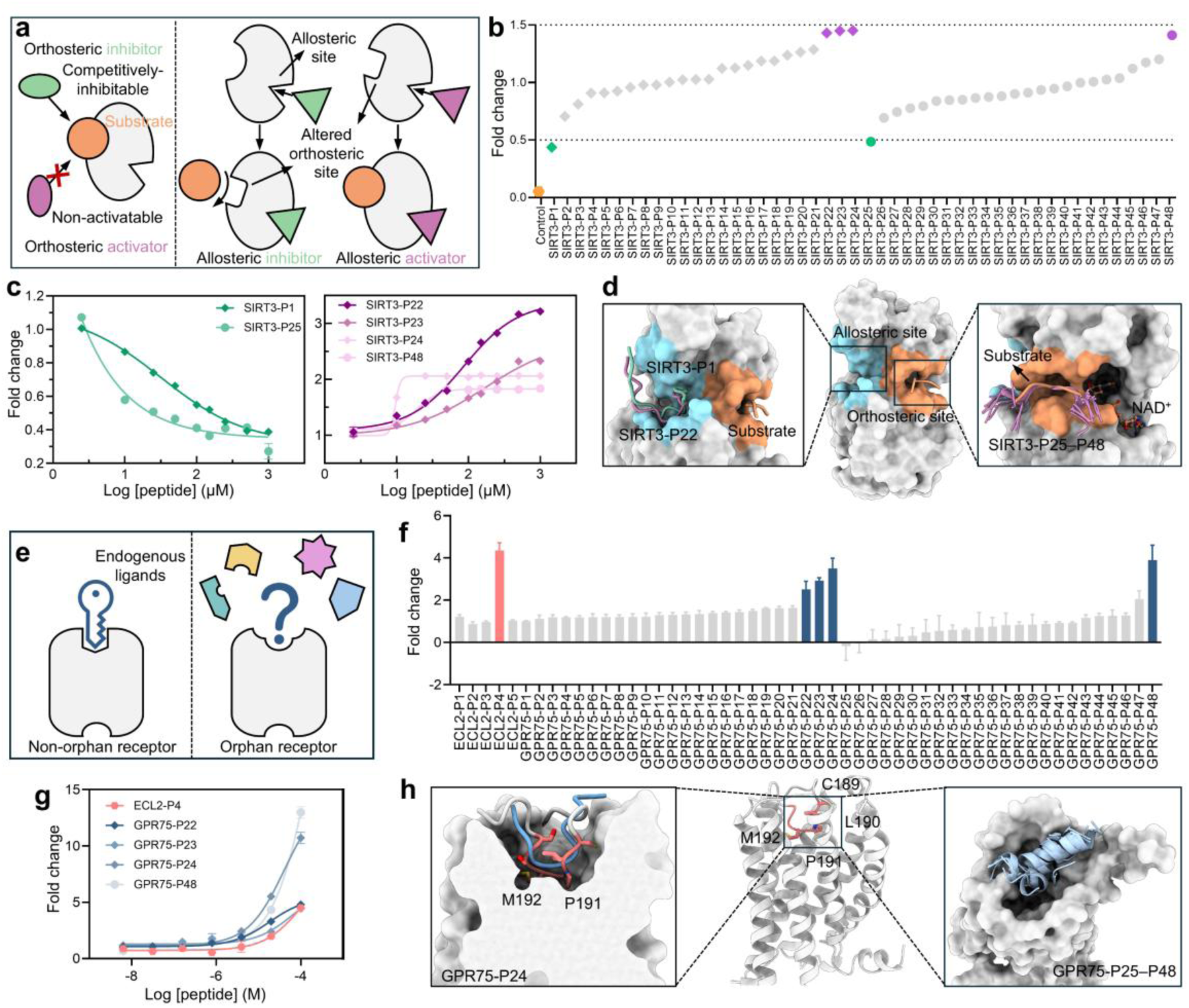
NeoPep-designed peptides targeting challenging undruggable targets. **a,** Schematic representation illustrating orthosteric versus allosteric regulation. Receptors are colored gray, substrates as orange circles, orthosteric agents as ellipses, and allosteric agents as triangles. Inhibitors and activators are colored green and purple, respectively. **b,** Functional screening of SIRT3 using a Fluor-de-Lys® (FdL)-based deacetylase activity assay. **c,** Dose–response curves for SIRT3 hits, displaying inhibitory (left, green) and agonistic (right, purple) activities. Data are shown as mean ± SEM (n = 3). **d,** Predicted binding poses of SIRT3-targeting peptides. SIRT3 (gray surface) with the allosteric site highlighted in cyan and the orthosteric site in orange. **e,** Schematic comparing non-orphan and orphan receptors. Endogenous ligands for non-orphan receptors are depicted as keys, and unknown ligands for orphan receptors as question marks surrounded by diverse candidate shapes. **f,** GPR75 activation screening measured via Tango reporter assay. Peptides exhibiting a fold change > 2 were classified as putative agonists. **g,** Dose–response curves for GPR75 agonists. The reference peptide ECL2-P4 is shown in red, and NeoPep-derived peptides are in blue. Data are presented as mean ± SEM (n = 3). **h,** Structural modelling of GPR75 agonistic peptides. GPR75 is shown in cartoon with ECL2 residues C189 to M192 displayed as sticks.

GPR75 is a class A GPCR without a known endogenous ligand, classified as an orphan receptor (Fig. 4e)^45,46^. Structural models from AF database indicate that GPR75 adopts an atypical architecture in which the putative orthosteric pocket is occluded by inward collapse of extracellular loop 2 (ECL2). This unusually buried ECL2 provides the structural basis for the constitutive activity of GPR75 (Supplementary Fig. 8). After we had concluded wet-lab experiments, a cryo-EM structure of apo GPR75 became available^47^. Although alignment of the AlphaFold-predicted structure with the cryo-EM model yielded a low overall RMSD of 2.0 Å, confirming the presence of the occluded pocket, the experimentally resolved backfolded ECL2 conformation differed markedly from the prediction (RMSD = 3.8 Å). This discrepancy highlights the difficulty of our initial de novo design, relying on a predictive structure with local inaccuracies. To test whether the ECL2 sequence contains an activating determinant, we firstly screened a series of truncated ECL2-derived peptides. Among them, ECL2-P4 elicited detectable receptor activation, with a potency of 102.4 μM and a span of 3.6-fold (Fig. 4f). In comparison, the AF2-multimer redocked cohort only yielded one notable candidate, GPR75-P48, which drove a higher span (12.3-fold) but diminished potency (EC_50_ = 174.0 μM). NeoPep-generated candidates consistently improved functional profiles. GPR75-P22 (EC_50_ = 18.4 μM) and GPR75-P23 (EC_50_ = 64.9 μM) maintained comparable agonistic response with greater potency, whereas GPR75-P24 achieved an optimally balanced profile, delivering a more than twofold enhancement in both potency (EC_50_ = 41.6 μM) and maximal response (span = 9.7-fold) over the derived ECL2-P4 baseline (Fig. 4g). Structural modeling implies that GPR75-P24 adopts a conformation resembling the native backfolded ECL2, providing a structural basis for its balanced activation (Fig. 4h, left). The high-confidence candidates predicted by AF2-multimer exhibited a systematic propensity to form α-helices (Fig. 4h, right). Indeed, we observed an increasing tendency to adopt helical conformations as AF2-multimer confidence increased, a trend also apparent in retrospective analysis of another GPCR, NTSR1 (Supplementary Fig. 9).

### Structure redesign for signaling selectivity

Building on NeoPep’s success in designing functional peptides for challenging targets, we next investigated whether its structure redesign mode could be leveraged to engineer signaling selectivity, an essential step toward uncoupling therapeutic efficacy from off-target toxicity. Such adverse effects typically arise either from activation of homologous receptor subtypes or distinct receptor conformations that trigger alternative signaling cascades^48^. The opioid receptors (ORs) represent a classic conundrum, as opioids remain highly effective analgesics^49^ yet are limited by severe side effects that underpin the ongoing opioid crisis^50^. Considerable effort has therefore been directed towards developing safer opioid ligands, including subtype-selective agonists (e.g., targeting κOR) and most notably, conformationally selective μOR agonists that preferentially engage G protein over β-arrestin^51^.

Initially, NeoPep was employed to design peptides against μOR in its G_i_-coupled conformation. This effort tested whether NeoPep could generate agonists for ORs, whose binding pockets are deeper and structurally distinct from those of previously studied GPCRs (Supplementary Fig. 10a), and whether a single-conformation design strategy could achieve selectivity over κOR and the β-arrestin-coupled state of μOR. Notably, two of the top-ranked candidates contained the N-terminal YGGF motif, the canonical minimal pharmacophore responsible for endogenous opioid recognition^52^. These two peptides, together with ten additional candidates, were synthesized and experimentally evaluated. Three peptides were confirmed as G_i_ agonism agonist of μOR in bioluminescence resonance energy transfer (BRET) assays, with EC_50_ values of 0.2 μM, 0.6 μM, and 22.7 μM, respectively (Supplementary Fig. 10b and c). We subsequently evaluated receptor subtype selectivity at κOR and conformational bias by measuring β-arrestin recruitment at μOR. All three agonists exhibited modest preference for μOR over κOR. Quantification of conformational bias using bias factors^53^ further revealed that these peptides elicited either balanced or only weakly biased signaling profiles (Supplementary Fig. 10d and e). Similarly, we measured β-arrestin recruitment for the six previously NeoPep-identified active NTSR1 peptides. These yielded bias factors ranging from 0.4 to 1.9, resulting in weak-to-moderate selectivity (Supplementary Fig. 11). To summarize, these findings indicate that NeoPep can identify μOR agonists within a dozen designs, yet a single-subtype-conformation design strategy remains insufficient for achieving profound subtype or conformational selectivity.

To achieve higher selectivity, we expanded NeoPep into a structure redesign framework. The pipeline begins with performing de novo sequence-and-structure co-design against a primary target state, either a specific receptor subtype or conformation state. Subsequently, for the alternative target state, the sequence module of NeoPep was held fixed while the structural module predicted the peptide binding pose. MD simulations were then performed on both poses to derive the difference in dynamic binding free energy (ΔΔG) (Fig. 5a and b). We first asked whether ΔΔG could distinguish the known preference of the YGGF tetrapeptide for μOR over κOR^52^. The predicted binding free energy for μOR (−45.0 kcal/mol) was 3.9 kcal/mol more favorable than that for κOR, supporting its validity. Thereupon, we deployed this workflow to design κOR-selective peptides over μOR. Of the 12 synthesized candidates, five showed higher κOR agonism in initial screening (Fig. 5c). Among them, the OR-P13 exemplified a clear κOR-selective profile with a potency of 3.5 μM and an efficacy of 90.2%, compared with 10.3 μM and 65.5% at μOR. The OR-P14 achieved even greater selectivity with an EC_50_ of 299.0 nM and an E_max_ of 81.7% at κOR, representing nearly 2,000-fold increase in potency relative to its marginal activity at μOR (EC_50_ = 595.8 μM; E_max_ = 39.8%) (Fig. 5d). The predicted structure pointed to two key features underlying the κOR selectivity of OR-P14. Its K^10^ establishes a tight salt bridge (2.7 Å) with D138^3.32^, mirroring κOR-selective small molecules JDTic (3.0 Å) and MP1104 (3.1 Å) but absent in the native peptide dynorphin^54^. In addition, the W^11^ engages a deep hydrophobic subpocket lined by Y320^7.43^ and W287^6.48^, resembling the spatial occupancy of the isopropyl and cyclopropylmethyl moieties in JDTic and MP1104 (Fig. 5e). These interactions are critical since disruption by residue mutation or cyclopropylmethyl truncation to methyl strongly attenuates potency^54^. This feature places OR-P14 within an emerging class of tryptophan-containing opioid peptides that achieve potent, subtype-selective agonism without the canonical tyramine pharmacophore^55^.

**Fig. 5.**
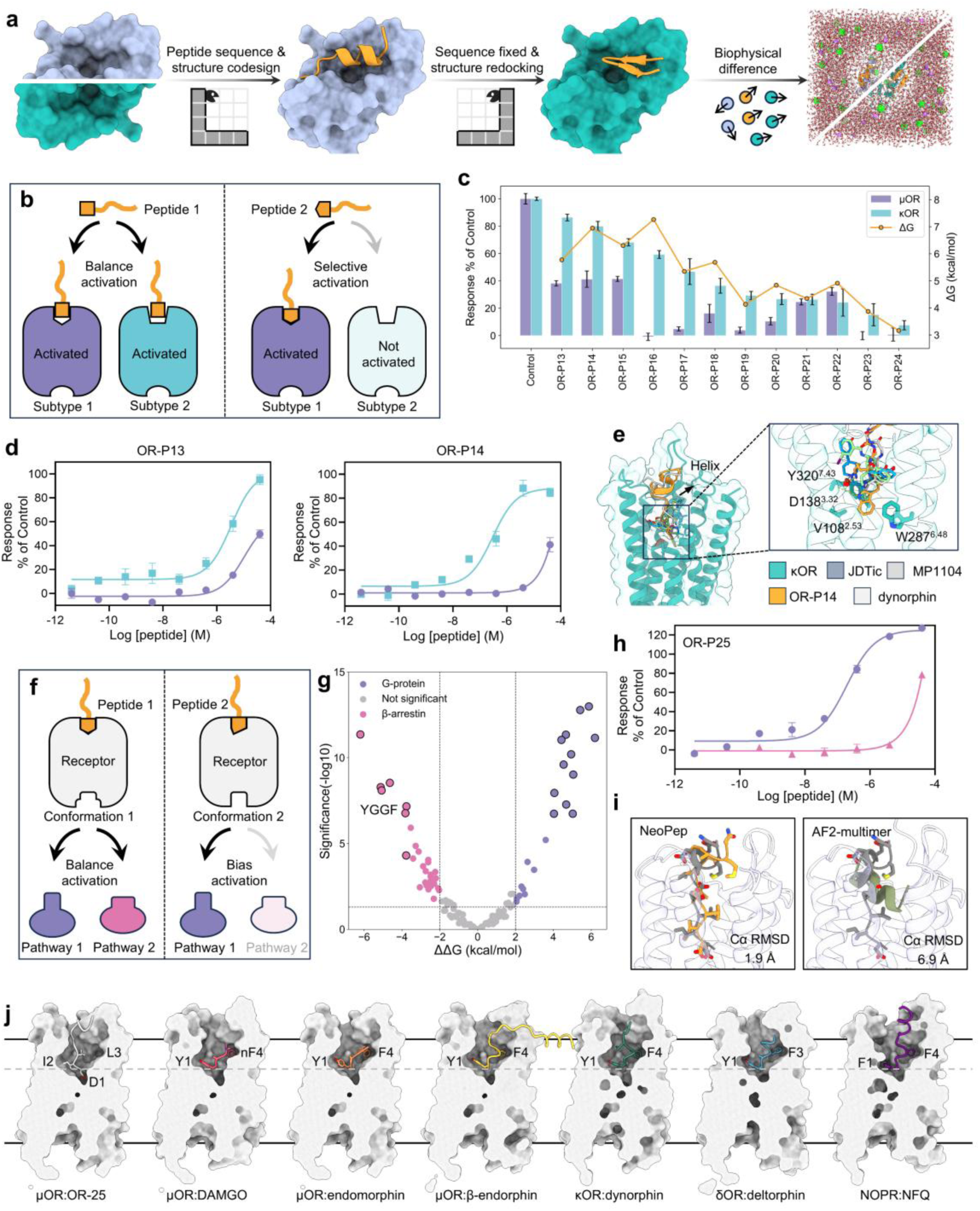
Rational design of selective peptides via NeoPep-driven structure redesign. **a,** Schematic of the NeoPep pipeline for peptide structure sampling. Serpentine representations in various orientations denote distinct peptide conformations of the same sequence. **b,** Schematic representation illustrating non-selective (balanced) versus subtype-selective receptor activation. Peptide 2 selectively activates subtype 1 (indicated by opaque coloring) over subtype 2 (indicated by transparent coloring). **c,** Functional screening of peptides across μOR and κOR. Bar chart displays the activation response for μOR (purple) and κOR (cyan), overlaid with the computationally predicted binding free energy (ΔΔG). Error bars represent SEM. **d,** Dose–response curves for OR-P13 and OR-P14, exhibiting κOR-selective profile (cyan) over μOR (purple). Data are mean ± SEM (n = 3). **e,** Superimposition of the OR-P14 modeled structure with cryo-EM structures of endogenous peptide dynorphin, small molecules JDTic and MP1104. Color usage: κOR, cyan; JDTic, blue; MP1104, green, OR-P14, orange and dynorphin, gray. **f,** Schematic comparing non-selective (balanced) with conformation-selective receptor activation. Peptide 2, when bound to receptor conformation 2, preferentially signals through pathway 1 relative to pathway 2. **g,** Volcano plot for pathway-selective peptide design. Variants are plotted by ΔΔG and statistical significance, separating substitutions favoring G-protein signaling (ΔΔG > 2 kcal/mol and *p* < 0.05, pink circle), β-arrestin signaling (ΔΔG < −2 kcal/mol and *p* < 0.05, purple circle), or no significant bias. The parental YGGF motif is indicated. Final synthesized candidates are highlighted by larger, black-bordered circles. **h,** Dose–response curves for OR-P25, demonstrating profound G-protein bias (purple) over β-arrestin recruitment (pink). Data are mean ± SEM (n = 3). **i,** Comparison of experimentally solved and computationally predicted binding modes of OR-P25. Left, cryo-EM structure of the OR-P25–μOR complex (grey) overlaid with the NeoPep design model (orange). Right, cryo-EM structure overlaid with the AF2-multimer predicted model (dark green). **j,** Cross-sectional surface representations opioid peptide binding poses. These structures compare of the designed peptide (OR-P25) against endogenous and synthetic reference ligands across the OR family (μOR, κOR, δOR, NOPR). The gray dashed line delineates a comparative reference plane for interaction depth.

We next programmed conformationally selective μOR agonists (Fig. 5f). To this end, we examined endomorphin binding in G_i_- and β-arrestin-coupled μOR complexes, computing a binding ΔΔG of −3.76 kcal/mol consistent with its known β-arrestin bias^52^. The overall ΔΔG distribution of generated peptides likewise suggested that β-arrestin-biased designs were more readily obtained than G_i_-biased counterparts (1.5×) (Fig. 5g). Nevertheless, given the greatest therapeutic potential of G_i_-biased μOR agonists to widen the therapeutic window and reduce adverse effects, we prioritized 12 top-ranked G_i_-biased candidates for experimental validation. An additional six β-arrestin-recruiting candidates were also included to demonstrate the bidirectional design versatility of our platform. As a result, OR-P25, OR-P26, and OR-P27 manifested G_i_-protein bias, with OR-P25 and OR-P27 achieving robust bias factors exceeding 100, whereas OR-P37 selectively engaged β-arrestin with a bias factor of 0.04 (Supplementary Fig. 12).

Remarkably, OR-P25 emerged as a potent, G_i_-selective agonist with an EC_50_ value of 194.6 nM, offering a promising structural lead for G_i_-selective μOR agonists (Fig. 5h). To validate the binding pose predicted by NeoPep and elucidate the molecular basis of the signaling bias, we solved the cryo-EM structure of the μOR–G_i_ complex bound to OR-P25 at overall resolution of 2.86 Å (Supplementary Fig. 13a and b, Materials and methods). The experimental structure closely matched the model generated by NeoPep, with a Cα RMSD of 1.9 Å and a heavy atom RMSD of 2.8 Å. By contrast, the top-ranked AF2-multimer model from extensive sampling adopted a helical conformation, a common artefact observed across the GPCRs we tested, and diverged markedly from the experimental structure (Cα RMSD = 6.9 Å and heavy atom RMSD = 7.7 Å) (Fig. 5i). The same was true for the best AF2-sampled pose irrespective of ranking, as well as for AF3 (Supplementary Fig. 13c). AF’s predictive inaccuracy probably stems from the atypical structural architecture of OR-P25 relative to known opioid-receptor templates. Unlike canonical opioid peptides^52^, OR-P25 lacks the conserved phenylalanine anchor, loosely engages the TM2-TM3 interface, and its residue D^1^ extends deeper into the TM core (Fig. 5j). A similar deep extension has been reported for the *N*-aniline and pyridine moieties of fentanyl and TRV130, respectively, and has been implicated in signaling bias^56^ (Supplementary Fig. 13d). Accordingly, an 18.2° tilt of the TRV130 pyridine ring relative to the *N*-aniline group of fentanyl weakens TM6/7 interactions, thereby attenuating β-arrestin recruitment (Supplementary Fig. 13e). The more pronounced 29.3° tilt of OR-P25 may underlie its exceptional G_i_ selectivity.

### Sequence redesign for functional refinement

This cryo-EM–validated atomic accuracy enables a further capability, iterative functional refinement, which we reformulate as a conditional inverse folding task over selected positions. In this framework, non-mutated residues are retained while targeted sites undergo redesign, allowing NeoPep to operate as a programmable mutator. In specific, the backbone structural module remains fixed at these designated positions, while the sequence output module predicts substituted amino acid (Fig. 6a). Following MM/PBSA calculation, the 12 variants with the most favorable binding energies are prioritized for experimental validation. Top-performing leads then serve as templates for the next optimization cycle. This iterative strategy mimics affinity maturation, empowering NeoPep to explore the local sequence space more efficiently than random mutagenesis while preserving the structural integrity of the binding interface. We validated this framework using the APC-inhibitory peptide described above. Although first-round de novo candidates displayed improved inhibitory activity relative to the derived scaffold, their potency remained in the high-micromolar range (IC_50_ = 28.6 μM), leaving room for further potency optimization.

**Fig. 6.**
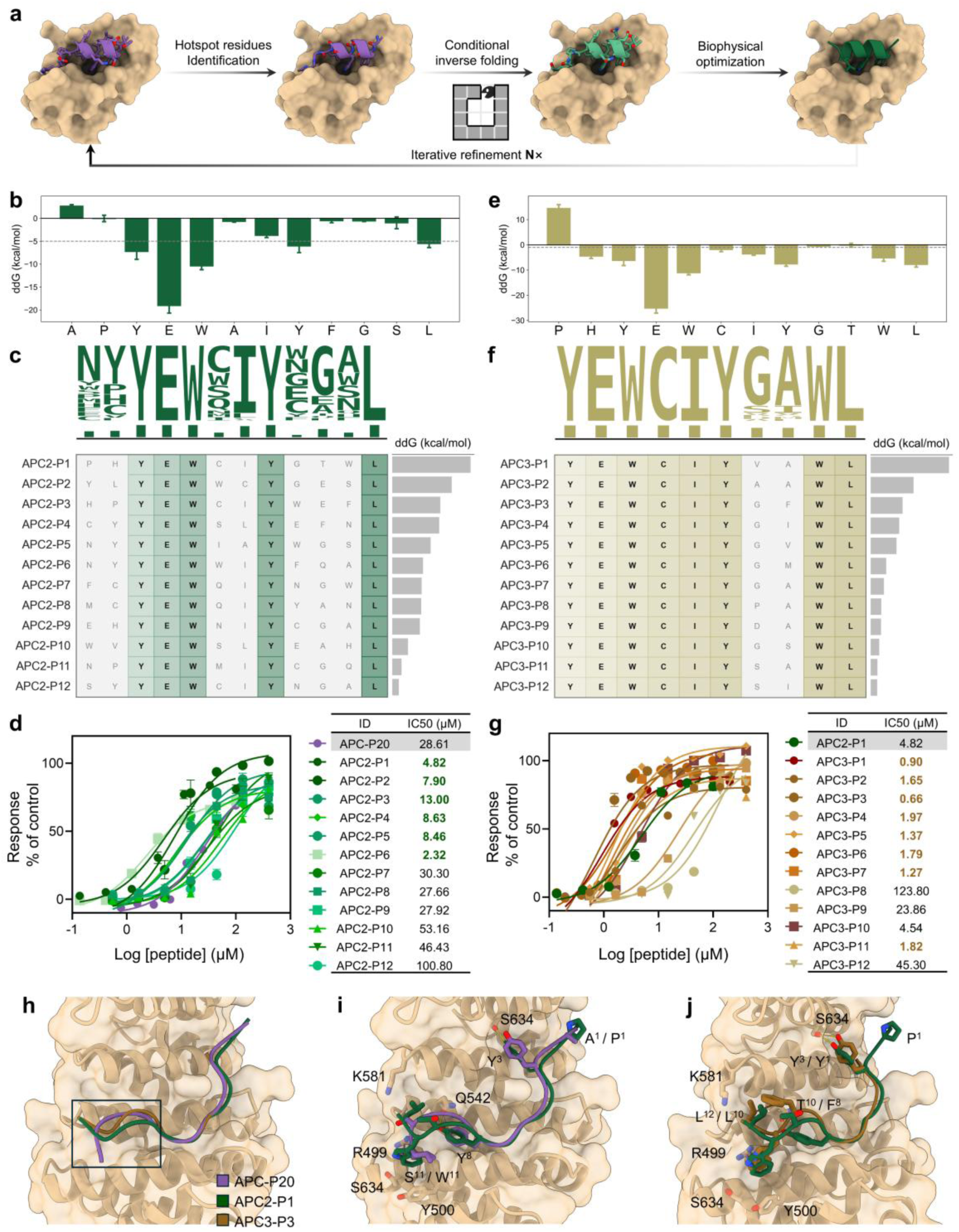
Rational design of enhanced APC peptides via NeoPep-mediated sequence redesign. **a,** Schematic of the NeoPep pipeline for peptide sequence evolution. The ouroboros representation marks iterative cycles of peptide refinement. **b, e,** Binding free-energy decomposition of the designed peptides. APC-P20 (**b,** green) and APC2-P1 (**e,** yellow). This color scheme applies throughout the figure. **c, f,** Sequence alignment of the top 12 peptides. Upper panels display sequence logos, where symbol height indicates residue frequency at each sequence position. Accompanying bar plots indicate the frequency of the most enriched residue per position. ΔΔG values, shown in the right panels, were calculated as the difference in ΔG between each peptide designed in the current round and the lead peptide from the previous round. **d, g,** Dose–response curves for the optimized peptides. Measured IC_50_ values are summarized in the adjacent tables, with high-potency variants highlighted in color. Data are presented as the mean ± SEM (n = 3). **h,** Structural superposition of the lead peptides from three successive rounds in complex with APC. **i, j,** Detailed analysis of binding modes across rounds. Key interactions are annotated: hydrogen bonds (yellow dashed lines), cation-π (purple lines), and hydrophobic contacts (gray).

In the second round, we incorporated MM/PBSA energy decomposition analysis to pinpoint residues with suboptimal binding contributions. Seven positions were selected for redesign under a threshold of −5 kcal/mol (Fig. 6b). We benchmarked NeoPep against the state-of-the-art sequence redesign model ProteinMPNN^57^, coupled with Rosetta for sidechain packing^30^, generating 100 unique variants for each method. ProteinMPNN manifested higher model confidence within a constrained sequence space, resulting in much lower sequence diversity than NeoPep. The 12 variants with lowest binding energy from each method were selected for experimental verification (Fig. 6c and Supplementary Fig. 14a). Moreover, six variants with the best ProteinMPNN scores were included for comparison (Supplementary Fig. 14b). As shown in Fig. 6d and Supplementary Table 3, 75% of the NeoPep-designed peptides produced improved or comparable potency relative to the first-round APC-P20, and half exhibited a marked improvement, with APC2-P1 and APC2-P6 achieving 5.9- to 12.3-fold enhancements in potency (IC_50_ = 4.8 and 2.32 μM, respectively). Conversely, ProteinMPNN-derived peptides yielded substantially inferior variants in both selection cohorts. None of the six top-ranked variants selected by ProteinMPNN score showed detectable activity. Prioritization by biophysical energy recovered weak inhibitory activity in five variants, with the best IC_50_ reaching 167.1 μM, but failed to salvage the optimization (Supplementary Fig. 14c and d).

A third iteration was performed using APC2-P1 as the template, selected over the comparably potent APC2-P6 due to its superior full inhibitory efficacy (97.8% vs. 81.8%). Reflecting a pharmacological preference for shorter, 10-amino-acid sequences, we truncated two N-terminal residues that energy decomposition analysis revealed to have negligible or unfavorable effects. Concurrently, half of the improved second-round variants retained I^7^, consistent with its marginal thermodynamic contributor (−3.9 kcal/mol) among the mutated sites. The redesign threshold was accordingly relaxed to −2 kcal/mol, and the top 12 peptides were taken forward for experimental validation (Fig. 6e), of which 75% exhibited stronger binding potency than their second-round predecessors (Fig. 6f, g and Supplementary Table 3). Although MD-guided assessment favored a Gly-Ala motif, driving a 3.8-fold improvement in potency, the NeoPep-generated candidates APC3-P1 and APC3-P3 achieved superior 5.4- to 7.3-fold enhancements, with APC3-P3 reaching an IC_50_ of 661.1 nM.

Collectively, two rounds of optimization culminated in a streamlined, 10-amino-acid peptide that delivered a 43.3-fold enhancement in potency. Structural modeling delineated that the N-terminal flanking residues (A^1^ in APC-P20 and H^1^ in APC2-P1) are dispensable for binding, whereas the C-terminal region underwent critical structural refinement (Fig. 6h). In APC2-P1, W^11^ forms a vital cation-π interaction with R499, anchoring the peptide in a lower subpocket defined by R499, Y500 and S634, while L^12^ engages an upper subpocket near Q542 and K581 (Fig. 6i). In the truncated APC3-P3, substitution of T^10^ to F^8^ likely stabilized this same C-terminal leucine (now L^10^) within the upper pocket, contributing to the observed gains in potency (Fig. 6j).

## Discussion

Peptides occupy a distinct biophysical niche, and discovery strategies optimized for small molecules or large biologics might not translate to functional peptide discovery^3^. In particular, accurately modelling the delicate and dynamic peptide–protein interactions de novo at this scale remains challenging for traditional physics-based approaches, whereas “black-box” deep learning models offer limited biophysical comprehension. NeoPep addresses these by advancing beyond implicit sequence pairing and purely geometric modelling toward robust sequence–structure–function mapping, anchored by over 5 million biophysically grounded data points. By integrating the efficiency of advanced deep learning with the interpretability of MD simulations, we could not only design programmable modes of peptide motion from scratch but also dissect atomic-level interaction networks underlying them.

In prospective validation across ten diverse and challenging targets, NeoPep achieved hit rates of 12.5–66.7%, with the top candidates exhibiting nanomolar potency. Such hit rates allow researchers to identify potent peptides by experimentally screening only dozens of designs, representing a substantial efficiency gain over random library-based screening, as demonstrated by a 50,813-fold enrichment for Trop2. Importantly, NeoPep can design peptides completely de novo from either a target structure or merely its primary sequence (coupled with AF monomer modeling, as in the case of GPC3 and GPR75), enabling the targeting of molecularly intractable proteins. Meanwhile, it not only generates high-affinity binders but also facilitates the bioengineering of imaging agents and therapeutic functional peptides, including inhibitors, activators, and allosteric regulators.

Benefiting from sequential fine-tuning on decoupled sequence and structure axes, NeoPep readily supports standalone structure and sequence redesign. It identified both μOR subtype-selective and G_i_ conformation-biased peptides. The cryo-EM structure of OR-P25-bound μOR confirms the structural precision of NeoPep (Cα RMSD = 1.9 Å), exceeding that of the best AF2 and AF3 predictions (Cα RMSD > 5.4 Å) under extensive sampling. Moreover, this atomic accuracy of NeoPep enables iterative sequence refinement without requiring resource-intensive experimental structures. This is evidenced by a 43.3-fold potency enhancement of an APC inhibitory peptide across three optimization rounds, transforming a weak native-derived peptide (IC_50_ > 100 uM) into a new 661.1 nM inhibitor. In contrast, the state-of-the-art ProteinMPNN failed to yield any improvement, underscoring the biophysical divergence between peptides and larger proteins. Combining NeoPep with rational pharmacokinetic design principles will enable the generation of therapeutic peptides simultaneously optimized for target potency, cellular permeability or oral bioavailability.

A widely utilized feature of AF2-multimer is its ability to estimate the confidence of a predicted structure. Although its ipTM metric acts as a strong binary predictor of binding activity in PPIs and achieves some success in peptide design^26,58^, our systematic investigation across diverse targets reveals that this proxy metric exhibits limited enrichment for positive functional peptides. The primary reason might be that such static metrics do not explicitly consider the thermodynamics of peptide binding, ignoring contributions from solvation and conformational heterogeneity. One notable observation is the strong correspondence between the NeoPep generations, MD simulations and experimental data. We attribute this agreement, at least in part, to biophysics-informed features of our approach, in particular the coordinated search across sequence and structure axes that progressively narrows to the designable seed anchor, although NeoPep does not explicitly model continuous conformational dynamics. In this respect, NeoPep and MD simulations are complementary.

Therefore, rather than rendering MD obsolete, we anticipate that generative frameworks like NeoPep will elevate accessible MD into an indispensable validation tool. A comparable paradigm shift is already underway with deep-learning force fields^59,60^. These emerging models aim to rapidly and accurately predict in silico conformational ensembles, an outstanding challenge for binary complexes, and particularly for transient peptide–protein complexes. Future development of deep learning integrated with the first-of-its-kind Dynamic dataset presented here could enable conformational ensemble generation and one-shot confidence prediction for the broader deep-learning community. Within the MD community, multiple kinetic models have been proposed to guide simulations via active learning loops, and an analogous strategy could be coupled with NeoPep to iteratively request new MD data targeted at minimizing uncertainty and increasing model confidence^61,62^.

We have demonstrated that by employing training-free guided flow matching under an optimal control framework, the pretrained model efficiently generates peptide mimetics directed by targeted mimicry rewards. Subsequent fine-tuning on these generated structures allows the model to internalize sequence–energetics mappings. In a similar vein, distance-constraint rewards can be seamlessly implemented for the zero-shot generation or iterative fine-tuning of conformationally restricted peptides^63^. Furthermore, by harnessing recent advances in all-atom diffusion and flow-matching models, we aim to extend this approach to the design of peptides incorporating non-canonical amino acids^64,65^.

## Materials and methods

### *In vivo* imaging of Trop2-targeting peptide tracers

BALB/c nude mice (female, 4-6 weeks old, HFK Bioscience) were raised under specific pathogen-free (SPF) conditions. The housing environment was regulated at 20–26 °C (with maximal temperature fluctuations of ≤ 3 °C) and 50–60% relative humidity, under a standard 12-h light/dark cycle. SKBR3 and A549 cells (1 × 10⁶ cells each) were suspended in a Matrigel-PBS mixture (1:1, v/v) and subcutaneously inoculated into the right axilla of BALB/c nude mice to establish tumor-bearing mouse models. Imaging commenced once tumor volumes reached approximately 150 mm³. The peptide was first conjugated to the bifunctional chelator DOTA. For radiolabeling, ^68^GaCl_3_ was eluted from a ^68^Ge/^68^Ga generator using 0.1 M HCl, adjusted with NaOAc buffer (pH 4.5), and incubated with DOTA-peptide conjugate at 95 °C for 15 min. The ^68^Ga-labeled product was purified via C18 solid-phase extraction (SPE). Tumor-bearing mice received an intravenous tail-vein injection of the ^68^Ga-labeled peptide (∼7.4 MBq per mouse, n=3). For competitive blocking experiments, 50 µg of unlabeled peptide was co-administered with the radiotracer. Micro-PET/CT scans were acquired at 15, 30, 45 and 60 min post-injection using a nanoPET/CT scanner (Mediso). Images were reconstructed applying a 3D ordered-subset expectation maximization (OSEM) algorithm. Regions of interest were delineated using InterView FUSION software, with tracer uptake across tumors and organs quantified as standardized uptake values. All animal experiments were conducted in accordance with the guidelines approved by the Institutional Animal Care and Use Committee of Fujian Medical University (IACUC approval, FJMU 2026-0048). Animals were euthanized at the conclusion of each experiment using approved humane endpoints.

### Immunohistochemistry

Mice were euthanized, and fresh tumor tissues were excised and immediately fixed in 4% paraformaldehyde solution. Tissues were embedded in paraffin, sectioned, and mounted onto poly-L-lysine-coated glass slides, then baked overnight at 60 °C to promote tissue adhesion. Sections were deparaffinized and rehydrated through a graded series comprising xylene (30 min), anhydrous ethanol (6 min), 95% ethanol (3 min), and 85% ethanol (3 min), and finally rinsed with distilled water for 1 minute. Antigen retrieval and endogenous peroxidase blocking were then performed, after which sections were blocked with normal goat serum for 10 min at room temperature. Sections were incubated with the primary antibody (anti-Trop2, EPR20043, Abcam, ab214488) for 60 min at room temperature, followed by the secondary antibody (goat anti-rabbit IgG H&L (HRP), Abcam, ab205718) in a humidified chamber for 20 min at room temperature. Color development was performed using freshly prepared 3,3’-diaminobenzidine (DAB) substrate solution, and sections were counterstained with hematoxylin for 2 min. After rinsing, the sections were mounted in neutral resin and examined by light microscopy.

### FdL-based deacetylase activity assay for SIRT3

Human SIRT3 (residues 118–399) was subcloned into a pET-28a-based construct with an N-terminal hexahistidine (His6) tag and expressed in Escherichia coli. SIRT3 protein expression and purification followed the well-established protocols^66^. Protein expression was induced with 0.5 mM isopropyl β-D-1-thiogalactopyranoside (IPTG; Sigma, I5502). Harvested cells were centrifuged, and the recombinant protein was isolated via Ni-NTA affinity purification.

FdL assays were performed to evaluate sirtuin activity as previously described^44^. Briefly, assays were carried out containing 250 nM SIRT3 protein, 10 μM substrate peptide (Z-(Ac) Lys-AMC), and 1 mM NAD⁺ (Sigma, V900401) in the absence or presence of test peptides at a final DMSO concentration of 1% (v/v). Reactions were incubated at 37 °C for 30 min, followed by termination via the addition of 10 μM Trypsin (Sigma, T4799) and 2 mM nicotinamide (Sigma, 72340). Fluorescence signals were recorded on a Synergy Neo microplate reader with excitation at 360 nm and emission at 460 nm.

### Tango reporter assay for GPR75

The Tango reporter assay was adopted as the primary functional readout for GPR75 activity measurements in this study. HTL cells were cultured in Dulbecco’s Modified Eagle Medium (DMEM) supplemented with 10% (v/v) FBS and seeded into 6-cm dishes (1.5 × 10⁶ cells/dish). After 18–24 h (70–80% confluency), cells were co-transfected with GPR75 and β-arrestin2-TEV constructs (1:1 DNA ratio, 1 μg each per dish). DNA was complexed with polyethylenimine (PEI; 2μ L/μg DNA) in Opti-MEM (Gibco) at a final DNA concentration of 10 ng/μL and incubated for 15 min at room temperature prior to cell addition.

Tango reporter assays were performed in white opaque-bottom 96-well plates (Beyotime, FCP968-80pcs). The following day, transfected cells were trypsinized, counted, and reseeded at 3.0–5.0 × 10⁴ cells/well in an 80-μL final volume. Test peptides were serially diluted in assay buffer to 5× working concentrations. Assays were carried out by adding the diluted peptides directly to the cells, followed by incubation for ≥16 h at 37 °C. The medium was subsequently aspirated and replaced with 100 μL of D-luciferin solution diluted 1:10 (v/v) in assay buffer (0.3 mg/ml). Following incubation for 6 min at room temperature in the dark, luminescence was recorded on an EnVision 2105 multimode reader (Revvity).

### BRET assays for NTSR1, μOR and κOR

HEK293T cells were cultured in DMEM supplemented with 10% (v/v) FBS. Cells were seeded into 10-cm dishes (3.0–4.0 × 10⁶ cells/dish) for NTSR1 assays, or 6-cm dishes (1.3–1.5 × 10⁶ cells/dish) for μOR and κOR assays. After 18–24 h (70–80% confluency), cells were co-transfected with specific plasmid combinations. For NTSR1 G_q_-protein dissociation assays, plasmids encoding NTSR1, Gα_q_–RLuc8, G_β_ and G_γ_–GFP2 were co-transfected (1:1:1:1 DNA ratio, 2 μg each per dish), while for NTSR1 β-arrestin2 recruitment assays, plasmids encoding β-arrestin2–GFP2, NTSR1–RLuc8 and GRK2 were co-transfected (10:1:1 DNA ratio, 8 μg total DNA per dish). For μOR and κOR G_i_-protein dissociation assays, plasmids encoding the receptor, Gα_i_–RLuc8, G_β_ and G_γ_–GFP2 were co-transfected (1:1:1:1 DNA ratio, 1 μg each per dish), while for μOR and κOR β-arrestin2 recruitment assays, plasmids encoding β-arrestin2–GFP2, receptor–RLuc8 and GRK2 were co-transfected (10:1:1 DNA ratio, 300 ng each per dish). Plasmids encoding the TRUPATH heterotrimeric G-protein complex were a gift from Bryan Roth (Addgene, kit #1000000163). DNA was complexed in Opti-MEM (Gibco) at a final DNA concentration of 10 ng/μL using either ExFect Transfection Reagent (Vazyme; 2 μL/μg DNA) for NTSR1 assays, or PEI (2 μL/μg DNA) for μOR and κOR assays.

BRET assays were performed in white opaque-bottom 96-well plates (Beyotime, FCP968-80pcs). The next day, transfected cells were reseeded at 3.0–5.0 × 10⁴ cells/well. For all assays, cells were stimulated with 20 μL of serially diluted test peptides in assay buffer (HBSS, 20 mM HEPES, pH 7.4) for 5 min, followed by the addition of 20 μL of freshly prepared coelenterazine 400a (5 μM). After a 2-min equilibration at room temperature, luminescence signals were recorded using either a Synergy Neo reader (BioTek) for NTSR1 or an EnVision multimode reader (Revvity) for μOR and κOR, both equipped with 410-nm and 515-nm emission filters. Then, BRET2 ratios were calculated by dividing the emission intensity at 515 nm by that at 410 nm. Signals were normalized to reference ligands (NTS^8–13^ for NTSR1, DAMGO for μOR, and dynorphin A^1–13^ for κOR), with the baseline set to 0% and the maximal reference response set to 100%.

### Fluorescence polarization (FP)-based competitive assay for APC

Human APC (residues 303–739) was subcloned into the pET-28a vector with an N-terminal His6 tag and expressed in *E. coli*. Cells were initially grown in Luria-Bertani (LB) broth containing kanamycin (50 μg/mL) for 6 h, then transferred to auto-induction ZYM-5052 medium containing kanamycin (100 μg/mL). Cultures were incubated at 37 °C for 6 h, followed by 18 h at 25 °C with shaking (250 rpm). Harvested cells (7,000 × g, 15 min) were resuspended in lysis buffer (25 mM Tris, 300 mM NaCl, 20 mM imidazole, 1 mM TCEP, pH 8.0) and disrupted using a high-pressure homogenizer. The clarified lysate (centrifuged at 23,000 rpm for 30 min) was applied to a HisTrap FF Ni-NTA column, washed with lysis buffer, and eluted with elution buffer (25 mM Tris, pH 8.0, 300 mM NaCl, 1 M imidazole, 1 mM TCEP). The eluate was dialyzed against storage buffer (50 mM HEPES, 300 mM NaCl, 1 mM EDTA, pH 7.5).

FP assays were performed in black 96-well plates (Corning, 3650). Recombinant APC was diluted in assay buffer (50 mM HEPES, 300 mM NaCl, 1 mM EDTA, 1 mM DTT, pH 7.5). Test peptides were prepared as concentrated DMSO stock solutions and serially diluted threefold in assay buffer. Assays were carried out in a final volume of 100 μL containing 27.4 nM APC protein, 20 nM FITC-labelled tracer peptide, and serially diluted test peptides. Test peptides were pre-incubated with APC protein for 1 h at room temperature, followed by addition of the FITC-labelled peptide tracer, and further incubation for 1.5 h at room temperature. Vehicle-treated wells and APC-free wells served as zero- and maximum-displacement controls, respectively. FP signals were recorded on a Synergy Neo microplate reader (BioTek) with excitation at 485 nm and emission at 528 nm.

### Cryo-EM data collection and structure determination

For cryo-EM grid preparation, 2.5 μL of the purified μOR-G_i_ complex bound to the de novo designed peptide agonist (5 mg/mL) was applied to glow-discharged holey carbon grids (Quantifoil R1.2/1.3, 300-mesh, Au). Grids were vitrified by plunge-freezing in liquid ethane using a Vitrobot Mark IV (Thermo Fisher Scientific) and stored in liquid nitrogen until data collection. Cryo-EM data were acquired at the Cryo-Electron Microscopy Research Center (Shanghai Institute of Materia Medica, Chinese Academy of Sciences) using a Titan Krios transmission electron microscope operating at 300 kV, equipped with a Falcon 4 direct electron detector. A total of 8,000 movies were recorded in electron event representation (EER) mode using EPU software at a calibrated pixel size of 0.73 Å and a defocus range of −0.8 to −1.8 μm. Each movie was acquired over 2.84 s with a total electron exposure of 50 e⁻/Å² and fractionated into 873 frames for motion correction.

Movie stacks were corrected for beam-induced motion using MotionCor v2.1. Contrast transfer function (CTF) parameters were estimated with CTFFIND4 within CryoSPARC v4. Particles were initially picked using the Blob Picker tool from micrographs with successfully determined CTF parameters and subjected to three rounds of 2D classification, yielding 1,549,708 initial particles. These particles were further processed by ab initio reconstruction followed by two rounds of heterogeneous refinement, resulting in a well-defined subset of 657,120 particles. Focused 3D classification, 3D Flex training, 3D Flex reconstruction, non-uniform refinement, and local refinement were subsequently performed in CryoSPARC v4, ultimately yielding a final reconstruction at an overall resolution of 2.86 Å from 80,000 particles.

For model building and refinement, the endomorphin-μOR-G_i_-scFv16 structure (PDB ID: 8F7R) was used as the initial reference model and rebuilt against the cryo-EM density map of the μOR-G_i_ complex^52^. The starting model was docked into the density map in UCSF ChimeraX and iteratively adjusted in Coot. Real-space and reciprocal-space refinement were subsequently carried out in PHENIX.

### Figure illustrations

Schematic illustrations were drafted in Microsoft PowerPoint. Functional study graphs were generated in GraphPad Prism (v10.4.2), structural figures were rendered in UCSF ChimeraX (v1.9) and Schrödinger PyMOL (v2.6), and Micro-PET/CT images were processed and displayed as described in the relevant methods section above. All remaining data plots were produced using Matplotlib.

## Supplementary Materials

Supplementary Methods 1 to 7

Supplementary Figures 1 to 14

Supplementary Tables 1 to 6

Supplementary References

